# Ray tracing models for estimating light collection properties of microstructured tapered optical fibers for optical neural interfaces

**DOI:** 10.1101/2020.05.07.083469

**Authors:** Emanuela Maglie, Marco Pisanello, Filippo Pisano, Antonio Balena, Marco Bianco, Barbara Spagnolo, Leonardo Sileo, Bernardo L. Sabatini, Massimo De Vittorio, Ferruccio Pisanello

## Abstract

Tapered optical fibers (TFs) were recently employed for depth-resolved monitoring of functional fluorescence in sub-cortical brain structures, enabling light collection from groups of a few cells through small optical windows located on the taper edge [1]. Here we present a numerical model to estimate light collection properties of microstructured TFs implanted in scattering brain tissue. Ray tracing coupled with *Henyey-Greenstein* scattering model enables the estimation of both light collection and fluorescence excitation fields in three dimensions, whose combination is employed to retrieve the volume of tissue probed by the device.

## MAIN TEXT

The possibility to genetically express fluorescent indicators of neural activity in sub-populations of neurons has made it possible to obtain unprecedented insight into signaling in the living brain. A wide set of molecular probes can be employed to monitor intracellular calcium transients [2], transmembrane potential [3] or neurotransmitter concentration [4]. These tools can be engineered to target specific cell types, and their overall response in terms of fluorescence dynamic range can be improved by restricting the expression specific subcellular compartments [3]. However, exciting and collecting fluorescence in sub-cortical structures of the mouse brain remains a challenge, as advanced multiphoton microscopy techniques are still limited to depths below ~1mm [5]. Fiber photometry (FP) is the most common method to monitor functional fluorescence in deeper brain regions [6]. In its most conventional implementation it exploits flat-cleaved optical fibers to excite fluorescence and collect the resulting time-varying optical signal. This method has enabled important applications to the study of functional connectivity [7], and it is compatible with fluorescence lifetime measurements [8], functional magnetic resonance imaging [9] and analysis in freely-moving animals and electrophysiology [10]. However, FP also shows important limitations when small cellular groups are targeted, since both excitation light and emitted fluorescence undergo tissue attenuation and scattering, strongly linking the detection volume size to the size of the fiber core [11].

Technologies have been proposed to overcome this limitation, including the use of implanted micro light emitting diodes (μLED) coupled to micro photodetectors (μPD) [12] and tapered optical fibers (TFs) [1,13]. With μLED/μPD pairs, the lower bound of detection volume is limited by the minimum size of the active elements and by the Lambertian emission profile of μLEDs. Conversely, TFs can be microstructured to tailor the volume over which signals are collected. This is achieved by covering the surface of the taper with a reflective metal to build a metal-confined narrowing waveguide, and then opening small windows, with Focused Ion Beam milling [14] or direct laser writing [15], through which light can couple to guided modes. However, the experimental estimation of emission and detection volumes from small detection points in living tissue is difficult, and, due to tissue heterogeneity, most certainly requires averaging across a high number of independent measurements. Conversely, although several works have proposed methods to estimate how light spreads in scattering tissue from implanted devices [16,17], the community would greatly benefit from generalized approaches to assess the collection efficiency of implanted photonic systems. We present a numerical method to evaluate the collection properties of optical windows on micro-structured TFs (μTFs), taking into account tissue attenuation and scattering. Collection efficiency maps are first evaluated experimentally in stained brain slices by using a two-photon microscopy system synchronized with light collection. These results are used as a point of comparison to implement a ray-tracing model based on a point source scanning in the volume close to the optical window, giving access to a three-dimensional (3D) collection efficiency map and to the related sensitivity volume. The combination of *collection efficiency maps η(x,y,z)* with emission patterns, obtained with the same ray-tracing approach, allows accessing the photometry efficiency field *ρ(x,y,z)*, thus estimating the collection properties when the same optical window is used to excite and collect fluorescence. μTFs were realized by micropatterning an aluminum-coated TF with FIB milling to open small apertures on the taper edge [14,18]. A ~600 nm-thick Al layer was deposited on a TF with numerical aperture (NA) 0.39, taper angle ψ=4°, and a 45 μm x 45 μm optical window was milled at a taper diameter of ~50 μm (**Fig. 1a**). The process was stopped when the entire thickness of the metal coating was removed.

**Fig. 1.**
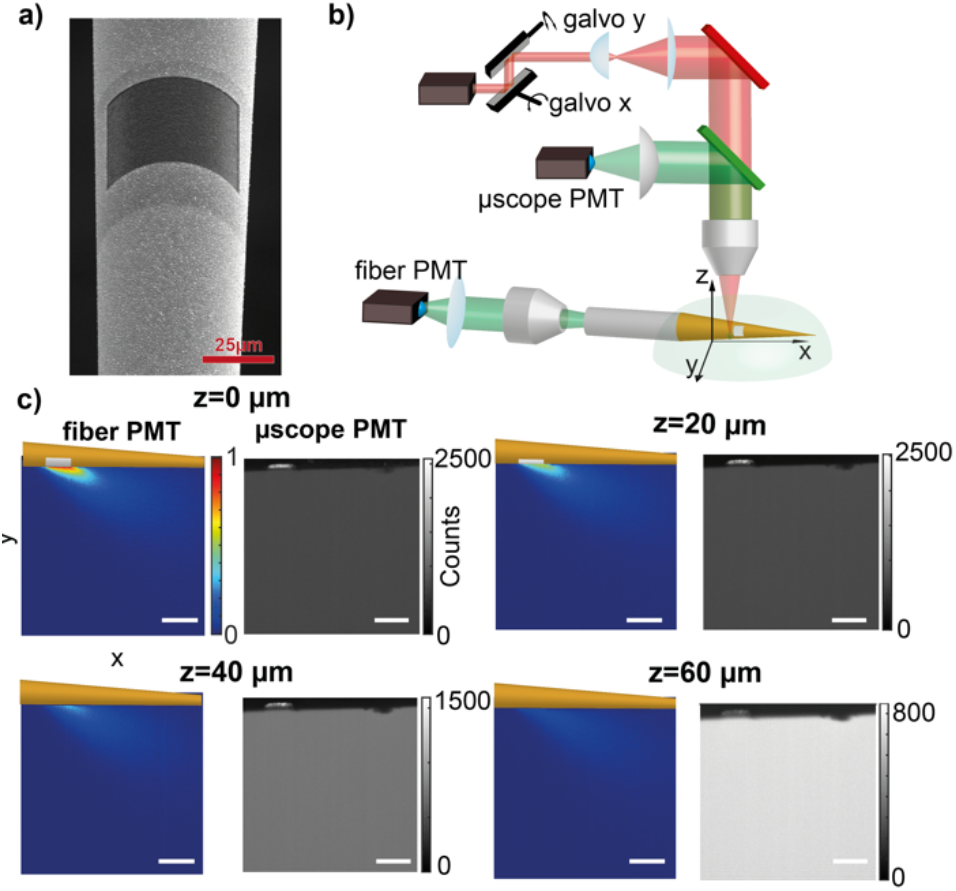
**(a)** Representative Scanning Electron Microscope (SEM) image of μTF with a 45 μm x45 μm optical window. (b) Optical setup implemented to measure the light collection properties of μTFs. (c) Representative data collected by a 45 μm x45 μm optical window and detected by the fiber PMT, with the related signal on the μscope PMT at different values of z.

Light collection properties of μTFs were first measured in a quasi-transparent phosphate saline buffer (PBS):fluorescein solution and then in fluorescein-stained coronal mouse brain slices to estimate the influence of tissue scattering. The optical setup implemented for this purpose is shown in **Fig. 1b** [11]. A fs-pulsed laser beam at λ=920nm was sent to a galvanometric mirrors (GMs) pair, a scan and tube lens with focal lengths of 50 mm and 200 mm, and an objective lens allowing for a FOV diameter of ~2.0 mm (Olympus XLFluor 4x/340a-NA=0.28). The fluorescence spot (in-plane point spread function 3 μm, axial size 32 μm [11]) was raster-scanned close to the optical window (**Fig. 1b**), with a fraction of the generated fluorescence signal collected by the optical window, propagated along the straight fiber and detected by a photomultiplier tube placed at the end of collection path (*fiber* PMT). Once synchronized to the GMs’ position, the data acquired by the *fiber* PMT provide the light collection field *η(x,y)* of the μTF under investigation. Part of the signal is also collected by the microscope objective and, through an epifluorescence path, it is sent to a second photomultiplier (*μscope* PMT) to estimate the uniformity of the generated fluorescence intensity in the FOV. Examples of the signals collected by the two PMTs are displayed in **Fig. 1c** for a μTF featuring a 45 x 45 μm optical window in a 30 μM PBS:fluorescein solution (for z=0 μm and z=20 μm, 40 μm, and 60 μm, respectively). Standard deviation of intensity detected by the *μscope* PMT in the scanning region was kept at ~4%. Similar considerations are valid also for data acquired at different positions along the z-axis, which give access to the 3D collection field *η(x,y,z)*, displayed for z=20, 40, and 60 μm in **Fig. 1c**.

The same μTF was tested in a 300 μm-thick coronal slice of mouse brain (**Fig. 2a**) uniformly stained with fluorescein. Slices were obtained from wild-type C57/Bl6 mouse fixed with 4% paraformaldeyde (PFA) and then incubated with Triton TX-100 for 30 min for permeabilization. The staining was performed in 1 mM PBS:fluorescein solution overnight. The resulting *η(x,y)* at z=0 μm is displayed in **Fig. 2b**. Due to tissue scattering, light collection is more confined close to the 45 x 45 μm optical window with respect to the measurements performed in quasi-transparent solution. This is confirmed by the comparison between the intensity decay along the collection lobe’s main axis in **Fig. 2c** (90% decay in brain and in quasi-transparent medium occurs after 90 and 150 μm, respectively).

**Fig. 2.**
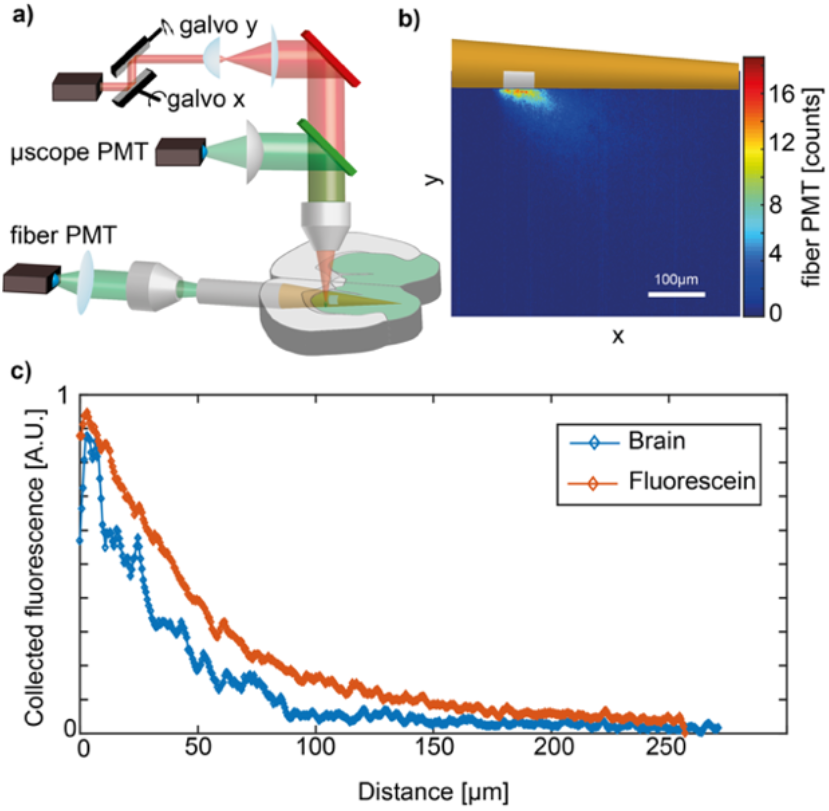
**(a)** Optical setup used for measurements in brain slices. (b) Representative data collected by the fiber PMT in brain slice. (c) Comparison between the intensity profile decay in brain slices (blue) and in PBS:Fluorescein solution (red).

The Ray Tracing Model (RT) was built using Zemax-Optic Studio (**Fig. 3a**), with a μTF featuring an optical window and a point source in the position (x, y) emitting at λ=520 nm. Rays emitted from the point source are scattered in the environment, enter the fiber through the window and are back-propagated toward a detector placed at the end of straight portion of the fiber of length l. To reconstruct η(x,y), each source position in the xy plane requires a RT run. A representative output of the simulations is displayed in **Fig. 3a**. The taper’s refractive index was set at n_t_=1.46 [19], while the surrounding environment at n_e_=1.36 approximating cerebral cortex’s refractive index [11]. The Henyey-Greenstein phase function was used to model brain scattering in agreement with recent literature, with photons mean free path μ_s_=0.04895 mm, anisotropy factor g = 0.9254 and transmission T = 0.9989 [11,16,20,21]. This model describes the angular distribution with which light rays are scattered:

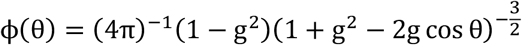

in which the anisotropy factor g defines if a scattering event is isotropic or anisotropic 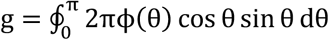.

**Fig. 3.**
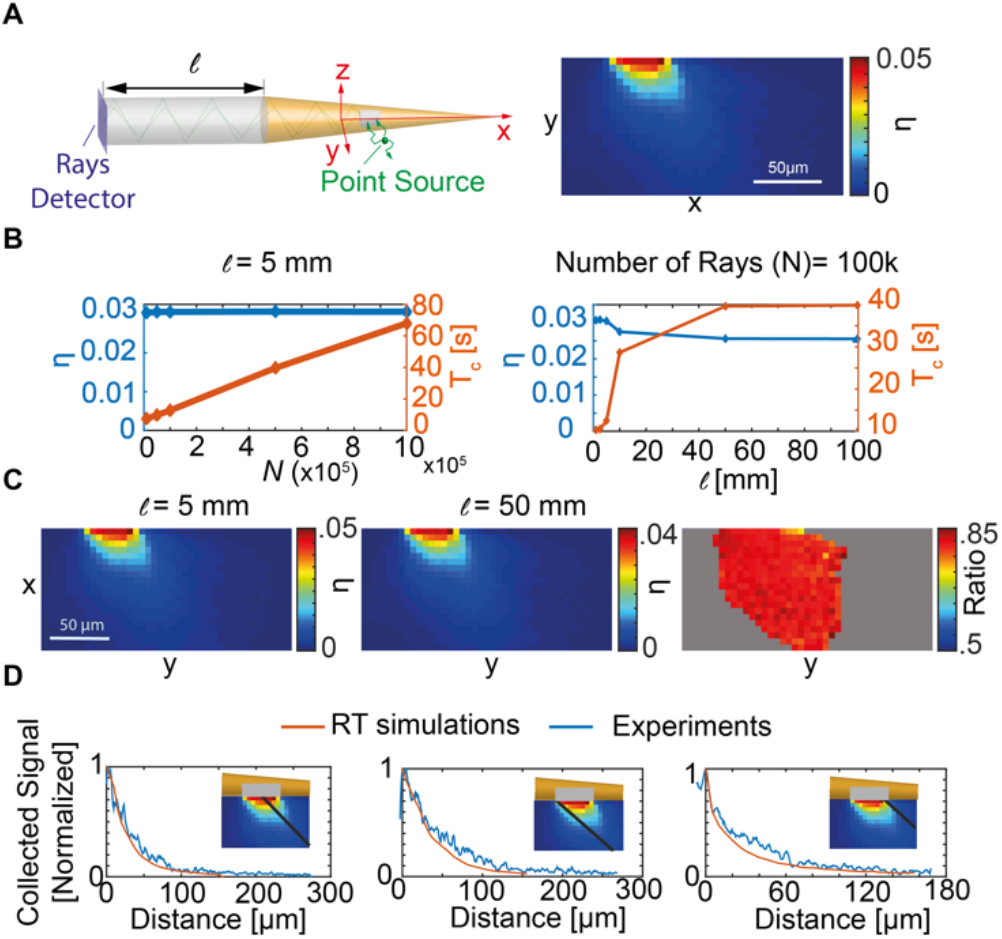
**(a)** Left: Schematic representation of the ray tracing layout used to estimate η. Right: representative η obtained by scanning the point source. (b) Convergence study as a function of l (Left) and N (Right). (c) Comparison between η(x,y) for two values of l (Left and Center), and their ratio (Right). The ratio is computed only were there a significative signal was found. (d) Comparison between experimental and numerical data along the profile lines shown in the inset.

The scattering event is assumed anisotropic and highly forward so that g approaches unity. Apart for the optical window area, the taper was coated by a 1 μm-thick Al layer set as ‘reflective’ surface with complex refractive index n_A1_=0.7-7.0i. The straight portion of the fiber was modeled in a step-index configuration with core/cladding diameters 200μm/225μm and refractive indexes of n_core_=1.4613 and n_clad_=1.4079, respectively. The length of the straight waveguide l and number of rays N used for each RT run were chosen after a convergence study (**Fig. 3b**), and set to the values of l=5 mm and N=10^5^ rays, allowing for a reasonable computational time. Convergence of the RT simulation was analyzed by running the model with the source at a fixed position while varying N or l. **Fig. 3b** left shows the variation of η and the computational time (T_c_) for N varying between 10^5^ and 10^6^ with l=5 mm. η remains constant ((3,00±0,01)×10^−2^), while T_c_ linearly increases. Therefore, N was set to 10^5^ in order to minimize computational times. **Fig. 3b** right shows the variation of η and T_c_ as a function of l in the range from l=1mm to l=100mm with N=10^5^. Both T_c_ and η converge to a plateau value starting from l=50 mm, with the single pixel simulation time T_c_ increasing from 5 s to 30 s for from l=5 mm to l=10 mm. We therefore investigated the differences in the simulation results for these two values of l, reported in **Fig. 3c** (left and center). **Fig. 3c** right shows the ratio between the data for the two lengths: by varying the position of the source within the entire simulation plane, the collected light intensities are proportional, with an average ratio in the simulation domain of 0.84 ± 0.01 (mean ± std deviation). To contain simulation times without affecting the simulation outcome, in the following, RT results are normalized to the maximum collected signal for l = 5 mm. In these conditions, we obtained a total run time of ~1.7 h in the 60×20 pixels domain, compared to >10 h required for l>10 mm. The size of the simulation domain was established after preliminary runs to find the adequate dimensions for measuring the entire volume with non-negligible signal collection. The comparison between RT simulation and experimental data for a 45 x 45 μm window is displayed in **Fig. 3d**, showing numerical and measured decays along three different axes across the collection lobe, as shown in the insets. The good agreement between experiments and simulations suggests that the proposed model can be employed to estimate collection properties from μTFs.

Extending the scan of the point source to the z direction allows assessing the three-dimensional collection field η(x,y,z) for the window and evaluating the overall collection volume V as a function of window size and position along the taper (**Fig. 4a** Left). The volumetric domain was set to 200 x 100 x 120 μm with a source point scan step of 10 μm, and extended to 200 x 100 x 200 μm for the largest window size (65 x 65 μm). After each simulation, η(x,y,z) maps were processed to estimate the volume enclosed by the iso-intensity surfaces at 10%, 20%, 40%, 60% and 80% of the maximum collection intensity (**Fig. 4a** Right). This was done for three different square windows with side W=25, 45, and 65 μm placed at three different locations (L=300, 750, and 1200 μm from the tip) along a taper with angle 4°). As shown in **Fig. 4b**, V increases as a function of W for all the investigated L, following the increase of window’s size at fixed distance from the taper’s tip. Instead, **Fig. 4c** displays the estimated volumes as a function of L for the different values of W. For the two smallest windows the collection volumes remain mostly unchanged along the taper, whereas for W=65 μm an increase of about one order of magnitude was recorded for L increasing from 300 to 1200 μm. Although this approach allows estimating collection fields from optical windows, in a typical fiber photometry experiment functional fluorescence is excited through the same device. Therefore, the intensity of the photometry signal depends on the geometrical distribution of excitation light. As reported by previous works, this can be described by the photometry efficiency (ρ) parameter, obtained by multiplying the collection field η(x,y,z) with the emission field normalized at its maximum β(x,y,z) [11,22]. A RT model was implemented to model the light emission field from an optical window placed along the taper, exploiting a stack of detectors close to the window to record irradiance and obtain the related emission field β(x,y,z) (**Fig. 5a**). A representative result of β(x,y,z) for a window with W=45 μm and L=750μm is displayed in **Fig. 5b**, together with the related collection field η(x,y,z) and the resulting photometry efficiency field ρ(x,y,z). **Fig. 5c** reports the volumes enclosed in the iso-surfaces at 10, 20, 40, 60 and 80% of the maximum signal for η (x,y,z) and ρ(x,y,z). **Fig. 5c** reports the volumes enclosed in the iso-surfaces at 10, 20, 40, 60 and 80% of the maximum signal for η (x,y,z) and ρ(x,y,z), highlighting the combined effect of light delivery and collection efficiencies.

**Fig. 4.**
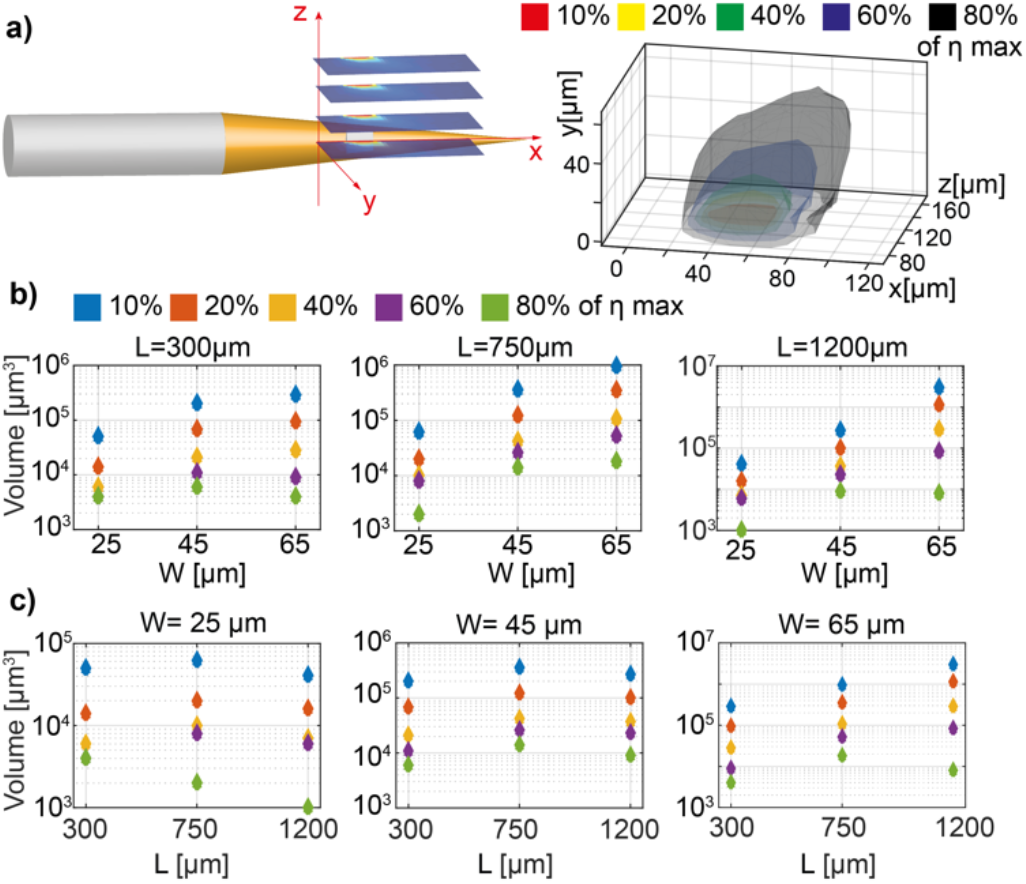
**(a)** Schematic representation of the 3D scanning (Left) and reconstructed iso-η surfaces used to estimate collection volumes. (b, c) Volumes contained within the iso-η surfaces as a function of W and 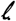

**Fig. 5.**
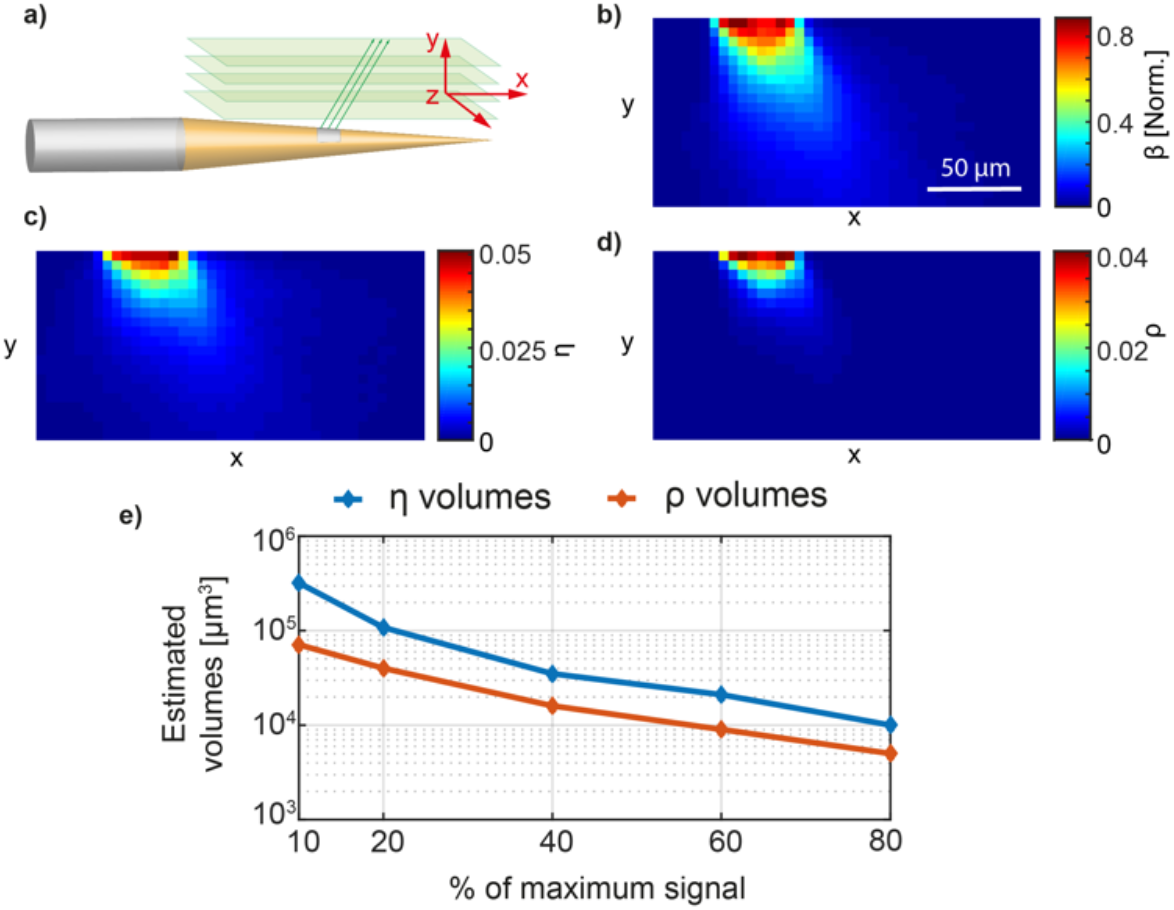
**(a)** Schematic representation of the ray tracing model used to estimate emission field. (b) Representative 2D emission field β(x,y) acquired at z=0 μm above the optical window. (c) η(x,y) on the same plane on the data in panel b. (d) Result of equation (1) applied to data in panels b and c. (e) Volumes contained in the iso-η and iso-ρ surfaces at different percentages of the maximum value.

In summary, we implement a RT method that estimates the collection properties of μTFs in living tissue, taking into account tissue attenuation and scattering. Our approach enables estimating the collection volume of optical windows with different sizes placed at different positions along the taper. In addition, the method can take into account the excitation efficiency resulting from the simulated emission pattern, in order to retrieve the photometry efficiency field, a foremost parameter in the design of fiber photometry experiments. Owing to the vibrant research surrounding the development of implantable neural probes [23] we view our method as a valuable resource for the community.

